# Interaction information along lifespan of the resting brain dynamics reveals a major redundant role of the default mode network

**DOI:** 10.1101/382705

**Authors:** Borja Camino-Pontes, Ibai Diez, Antonio Jimenez-Marin, Javier Rasero, Paolo Xu, Charles C.Y. Bonifazi, Sebastiano Stramaglia, Stephan Swinnen, Jesus M Cortes

## Abstract

Interaction Information generalizes the univariate Shannon Entropy to triplets of variables, allowing the detection of redundant or synergetic interactions in dynamical networks. Here, we calculated interaction information from functional magnetic resonance imaging and asked whether redundancy or synergy vary across brain regions and along lifespan. We found high overlapping between the pattern of high redundancy and the default mode network, and this occurred along lifespan. The pattern of high values of synergy, more heterogeneous and variable along lifespan, was overlapping with different cognitive domains, such as spatial and temporal memory, emotion processing and motor skills. Moreover, the amount of redundancy and synergy seem to be balanced along lifespan, suggesting informational compensatory mechanisms in brain networks.

## Introduction

Interaction information (II) quantifies in triplets of variables the amount of redundant (positive interaction) or synergetic (negative interaction) information contained in the triplet [1, 2]. While the mutual information (MI) shared between two variables is always positive or zero (for the case of independent variables), II can be either positive or negative, respectively, unveiling redundancy (R) or synergy (S). To give some specific examples, common-cause structures lead to R, whilst the combination of one XOR gate with two independent random inputs leads to S [3, 4].

The presence of synergetic effects is well-known to occur in sociological and psychological modeling, where (very often) there are some variables that increase the prediction power on different ones [5]. On the other hand, redundancy have been addressed before in gene regulatory networks [6, 7] and electrophysiological data in patients with epilepsy [2] or with deficit of consciousness [8], but, the pattern of triplet interactions in functional magnetic resonance imaging is not yet well-understood. By using a different methodology based on Granger causality influence, the authors in [9] found that R regions occurred mainly due to voxel-contiguity and inter-hemispheric symmetry, while S occurred mainly between non-homologous region pairs in contra-lateral hemispheres.

The functional connectivity (FC) patterns at rest have been shown to be altered in different pathological conditions such as deficit of consciousness [10, 11, 12, 13], schizophrenia [14, 15], epilepsy [16] and Alzheimer’s Disease [17, 18, 19, 20, 21]. Here, following a recent study [22] combining functional and structural multi-scale connectivity along lifespan, we address how redundancy and synergy varies from young to old people, within an age interval of 10 to 80 years old, that as far as we know, has not been addressed before.

## Material & Methods

### Participants

Participants were recruited in the vicinity of Leuven and Hasselt (Belgium) from the general population by advertisements on websites, announcements at meetings and provision of flyers at visits of organizations, and public gatherings (PI: Stephan Swinnen). A sample of *N* = 164 healthy volunteers (81 females) ranging in age from 10 to 80 years (mean age 44.4 years, SD 22.1 years) participated in the study. All participants were right-handed, as verified by the Edinburgh Handedness Inventory. None of the participants had a history of ophthalmological, neurological, psychiatric, or cardiovascular diseases potentially influencing imaging or clinical measures. Informed consent was obtained before testing. The study was approved by the local ethics committee for biomedical research, and was performed in accordance with the Declaration of Helsinki.

### Imaging acquisition

Image acquisition was performed in a magnetic resonance imaging (MRI) Siemens 3T MAGNETOM Trio MRI scanner with a 12-channel matrix head coil. The anatomical data was acquired as a high-resolution T1 image with a 3D magnetization prepared rapid acquisition gradient echo: repetition time (RT) = 2,300 ms, echo time (ET) = 2.98 ms, voxel size = 1 × 1 × 1.1 mm^3^, slice thickness = 1.1 mm, field of view = 256 × 240 mm^2^, 160 contiguous sagital slices covering the entire brain and brainstem.

Resting state functional data was acquired with a gradient echo-planar imaging sequence over a 10 min session using the following parameters: 200 whole-brain volumes with TR/TE = 3,000/30 ms, flip angle = 90, inter-slice gap = 0.28 mm, voxel size = 2.5 × 3 × 2.5 mm^3^, 80 × 80 matrix, slice thickness = 2.8 mm, 50 oblique axial slices, interleaved in descending order.

### Imaging preprocessing

We applied resting fMRI preprocessing similar to previous work ([23, 24, 25, 26, 27, 28]) using FSL and AFNI (http://afni.nimh.nih.gov/afni/). First, slice-time was applied to the fMRI data set. Next, each volume was aligned to the middle volume to correct for head motion artifacts. After intensity normalization, we regressed out the motion time courses, the average cerebrospinal fluid (CSF) signal and the average white matter signal. Next, a band pass filter was applied between 0.01 and 0.08 Hz [29]. Next, the preprocessed fuctional data was spatially normalized to the MNI152 brain template, with a voxel size of 3 x 3 x3 mm^3^. Next, all voxels were spatially smoothed with a 6 mm full width at half maximum isotropic Gaussian kernel. Finally, in addition to head motion correction, we performed scrubbing, by which time points with frame-wise displacements ¿0.5 were interpolated by a cubic spline [30]. We further removed the effect of head motion using the global frame displacements as a noninterest covariate, as old participants moved more than the young (when representing the mean frame-wise displacement as a function of age provided a correlation value equal to 0.51 with p value equal to 1e − 11), and this fact introduced nuisance correlations with age.

### Brain hierarchical atlas

The brain was divided in 2,514 brain regions that we grouped into modules using the brain hierarchical atlas (BHA), developed in [31] and applied by the authors in a traumatic injury study [32] and in a lifespan study [22]. The BHA is available to download at http://www.nitrc.org/projects/biocr_hcatlas/. A new Phyton version that was developed during Brainhack Global 2017-Bilbao can be download at https://github.com/compneurobilbao/bha.

Although full details have been provided before [31], very briefly, the use of the BHA guarantees three conditions simultaneously: (1) that the dynamics of voxels belonging to a same module is very similar, (2) that those voxels belonging to a same module are structurally wired by white matter tracts, (3) that modules are simultaneously functional and structural.

Here, we focus on the M=20 module partition as was shown to be optimal based on cross-modularity X [31], and index defined as the geometric mean between the modularity of the structural partition, the modularity of the functional partition, and the mean Sorensen similarity between modules existing in the two structural and functional partitions.

### Shannon entropy

The Shannon entropy of a random variable X is defined as:

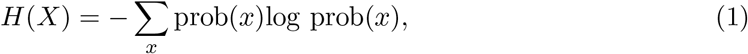

where *x* represents one possible state of variable *X* [33, 34]. Equation (1) can be generalized to two and three dimensions, respectively as *H* (*X, Y*) = − *Σ _x_ Σ_y_* prob(*x, y*)log prob(*x, y*) and *H*(*X,Y,Z*) = − *Σ_x_ Σ_y_ Σ_z_* prob(*x, y, z*)log prob(*x, y, z*). For base 2 logarithm (as we have done here), the entropy is expressed in bits.

Here, *X* represents any time series of resting state functional dynamics.

Interaction information

The interaction information (II) is an extension of the Shannon entropy to triplets of variables [1]. For any triplet (*x, y, z*), the interaction information (II) is defined as

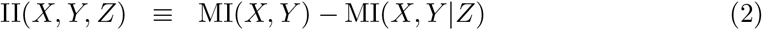

where MI(*x, y*) = *H*(*x, y*) — *H*(*X*) — *H*(*Y*) is the mutual information between *X* and *Y* and MI(*X, Y*|*Z*) is the conditional mutual information between *X* and *Y* conditioned to *Z* (for details see [34]).

The sign of II has important physical implications; when II is positive, the three variables (*x, y, z*) are said to be redundant, while if II is negative, the interaction in (*x, y, z*) is synergetic.

Here, *X, Y, Z* represent any three time series of resting state functional dynamics.

#### Calculation of II

II was calculated by applying equation (2) and estimating *MI*(*x, y*) and *MI*(*X, Y*|*Z*) using the Gaussian copula approach recently derived by [35]. In particular, we made use of the functions *cmL·ggg.m* and *mL·gg.m*; available at https://github.com/robince/gcmi/.

Important to remark is that because the copula entropy does not depend on the marginal distributions of the original variables, the authors in [35] transformed the marginals to be standard Gaussian variables, and therefore, the mutual information was exactly estimated under the Gaussian assumption.

#### Per module R and S

Values of R per brain module were obtained by summing (for a fixed module *m*) over all pairs such that II was positive, i.e.,

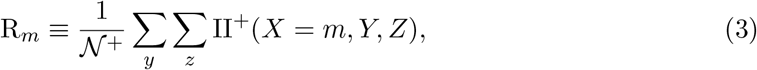

where II^+^ represent any positive value of II and *𝒮*^+^ the total number of positive elements. Analogously, the per module S was defined as:

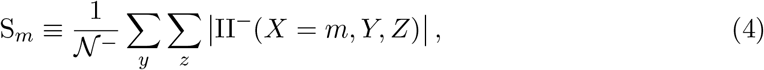

where II^−^ represent any negative value of II, *𝒮*^−^ the total number of negative elements and | · · · | absolute value.

For the calculation of both *R_m_* and *S_m_* we only considered triplets in which the three variables are distinct each other, ie. satisfying that *Y* ≠ *m*, *Y* ≠ *Z* and *Z* ≠ *y*.

Normalized values of R and S were calculated by dividing each value by its maximum.

### Mask of the resting state networks

Following [36], we created a mask for the different resting state networks by defining voxels with z-score value satisfying *Z* < −3 or *Z* > 3. In particular, we built masks for the default mode, cerebellum, executive control, frontoparietal, sensorimotor and visual resting state networks.

These masks were used to calculate the percentage of overlap between brain maps of R, S, 1 –S and R/S with the different functional resting state networks.

## Results

M=20 modules of the BHA were used as regions of interest. We calculated II for all possible triplets. Redundancy and synergy were assessed using equations (3) and (4) dividing the entire population (*N* = 164) in 4 different intervals of age: I1 (10-20 years old, *N*_1_ = 30), I2 (20-40 years old, *N*_2_ = 46), I3 (40-60 years old, *N*_3_ = 29) and I4 (60-80 years old, *N*_4_ = 59).

Values of R in bits are represented in figure 1. Panel a shows the values of R per each of the M=20 modules, at different age intervals. Panel b shows the average R across the M=20 modules. Significant statistical differences with age were found across the 4 groups (one-way ANOVA, p-value of p= 1e^−51^). Post-hoc analyses revealed statistical differences between all intervals with respect to I4, indicating an overall redundancy increment for the old population (I1 vs I4 p= 3e^−24^, I2 vs I4 p= 4e^−30^, I3 vs I4 p= 3e^−33^). All these comparison were significant after Bonferroni correction (i.e., using a thershold for significance of 0.05/6, being 6 the total number of pairwise comparisons). No other group comparison was significant after correction.

**Figure 1.**
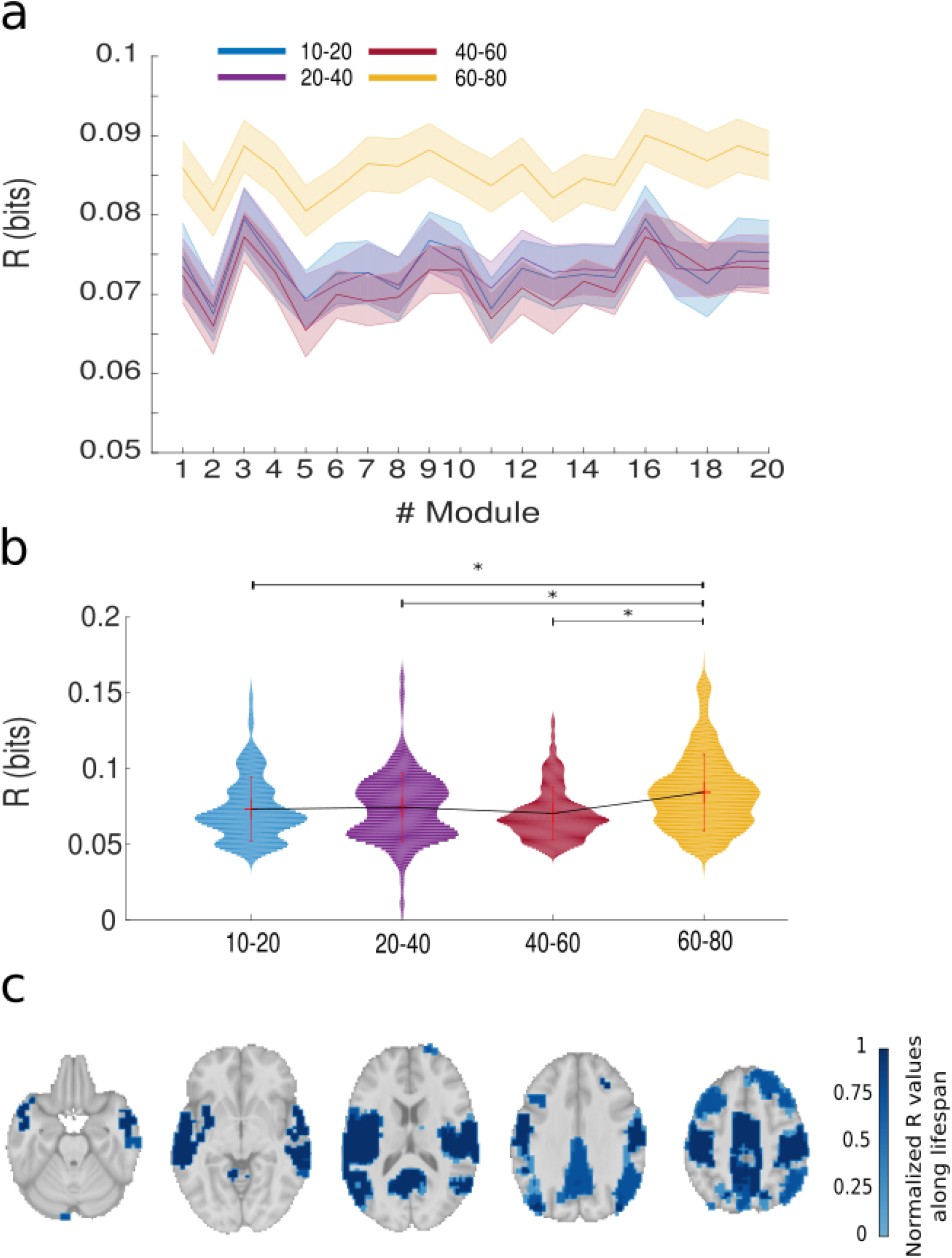
Across subjects, average redundancy (R) per brain module and along lifespan. **a:** For each module, values of R are represented in 4 different age intervals: blue (10-20 years old), purple (20-40), red (40-60) and magenta (60-80). Dark central lines represent average values across subjects and shaded areas represent statistical error **b:** Violin plots of R averaging over all modules and subjects within age interval. * represents statistical significant differences after Bonferroni correction. Notice that all age groups showed differences with respect to the old group (magenta), indicating that the overall R significantly increased in the old population. **c:** Brain maps of normalized R averaging over all age intervals with a threshold value of 0,7.

Brain maps of normalized R values per module are represented in figure 1c. Highest values were found in modules 3, 9 and 16, that bilaterally are located in cerebellum, precuneous, posterior cingulate, superior and middle temporal gyrus, paracentral lobule, precentral gyrus, superior frontal and parietal gyrus and insula. The function associated to these high redundant areas is a superposition of three important resting state networks, namely, default mode, sensory-motor and auditory networks (for further details see descriptive Table S1 in [31]).

Values of S in bits are represented in figure 2. Panel a shows the values of S per each of the M=20 modules at different age intervals. Panel b shows the average S across the M=20 modules. Significant statistical differences with age were found across the 4 groups (one-way ANOVA, p= 2e^−22^). Post-hoc analyses revealed significant differences for several comparisons after one-way anova (I1 vs I4 p= 2e^−18^, I1 vs I3 p= 3e^−5^, I2 vs I4 p= 5e^−15^, I3 vs I4 p= 6e^−6^). No other comparison survived after Bonferroni correction. Thus, in comparison to R, the dynamical pattern of S along lifespan is more heterogeneous.

**Figure 2:**
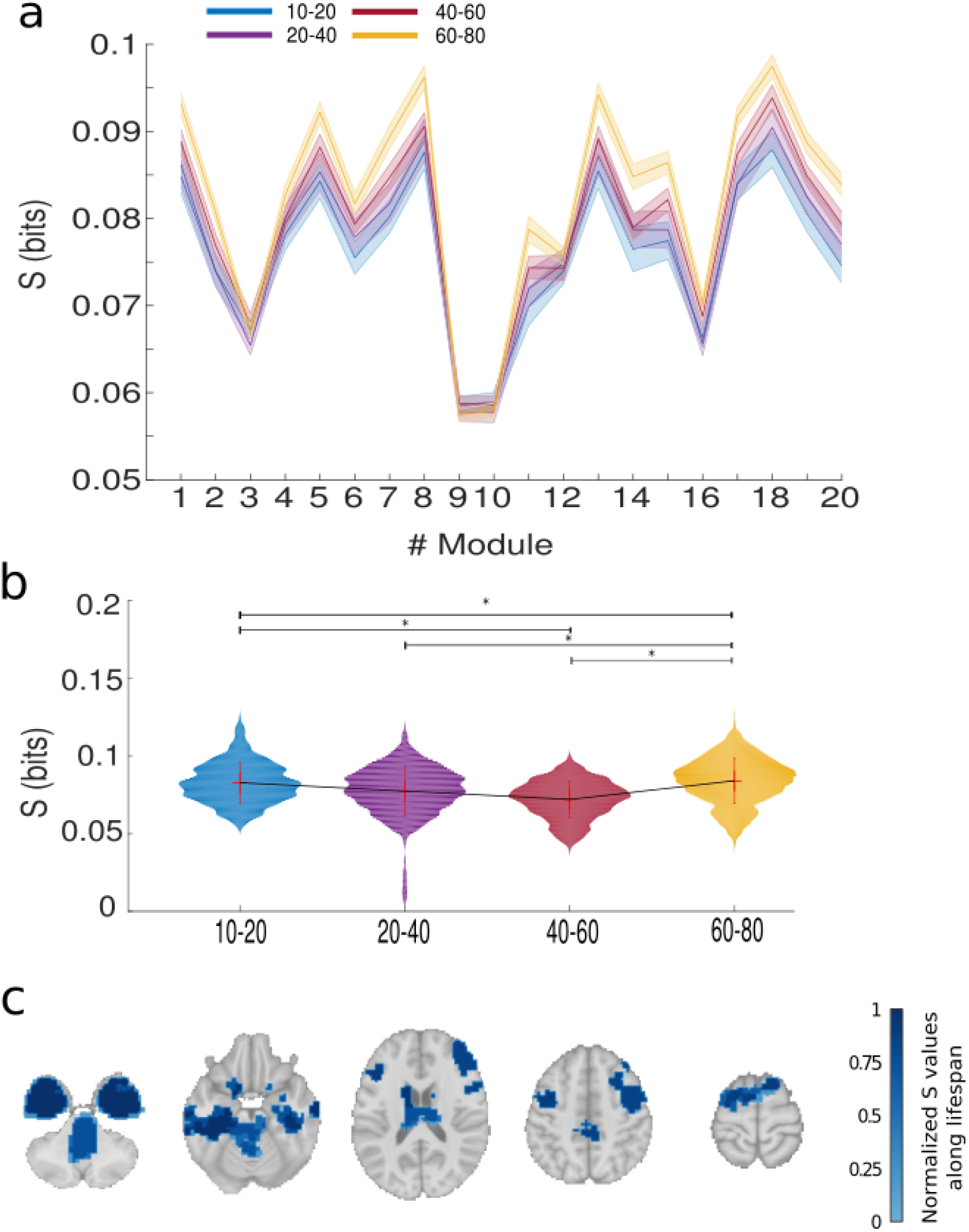
Across subjects, average Synergy (S) per brain module and along lifespan. **a:** For each module, values of average S are represented in 4 different age ranges, blue (10-20 years old), purple (20-40), red (40-60) and magenta (60-80). Dark central lines represent average values across subjects and shaded areas represent statistical error. **b:** Violin plots of S averaging over all modules and subjects within age interval. * represents statistical significant differences after Bonferroni correction. **c:** Brain maps of normalized S averaging over all age intervals with a threshold value of 0,7.

Brain maps of normalized S values per module are represented in figure 2c. Highest values were found in modules 3, 8 and 18, that bilaterally are located in hippocampus, amygdala, entorhinal cortex, fusiform, temporal pole, inferior temporal gyrus, caudate and putamen. These areas are associated to different cognitive domains, such as spatial and temporal memory, emotion processing and motor skills.

Although in general, by construction of R and S, the two measures are inversely proportional one to another, the two measures are in fact complementary. Figure 3 shows brain maps of normalized R together with the ones for 1–S. In particular, values with highest 1–S were found in modules 3, 9 and 10, bilaterally located in the anterior cingulate, inferior parietal and frontal gyrus, orbital gyrus, pars opercularis, pars orbitalis, pars triangularis, paracentral lobule, precentral gyrus, postcentral gyrus, precuneus, superior temporal gyrus, insula, cerebellum, posterior cingulate, inferior parietal gyrus, superior frontal gyrus. Therefore, brain maps of 1–S incorporate the frontal pole, increasing the overlapping with the default mode network from 50.32 for R (figure 3a) to 66.95% for 1–S (figure 3b).

**Figure 3:**
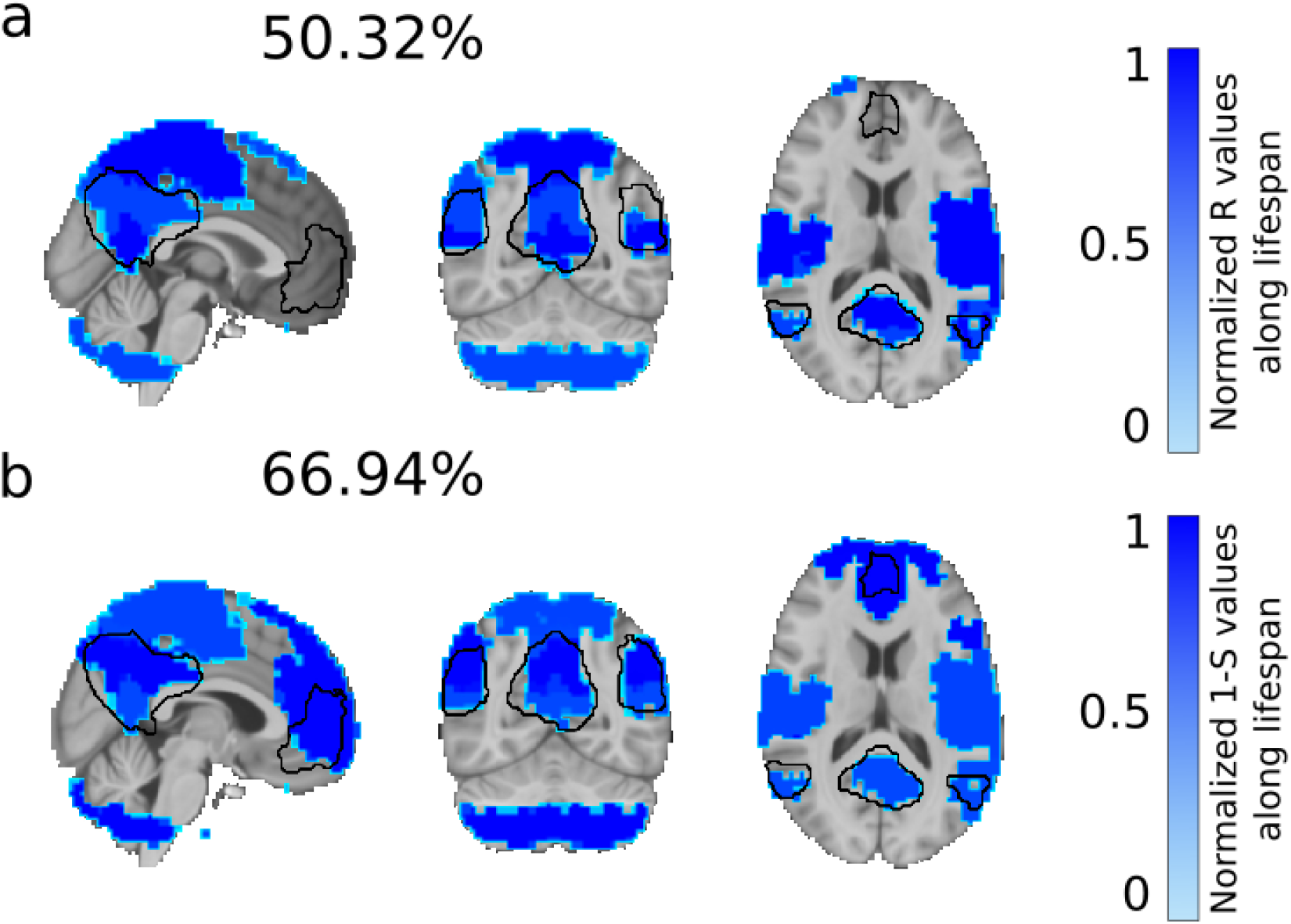
Normalized values of R and 1-S across brain regions reveals a key redundant role of the default mode network. **a:** Brain maps of normalized R averaged over age intervals, subjects and modules overlap 50.32% with a mask of the default mode network (depicted in black). **b:** Similar to panel a, but plotting brain maps of 1-S provided an overlap with the default mode network of 66.94%. Notice that 1-S but not R incorporates the frontal pole into the brain map, increasing the matching with the default mode network.

The amount of R is somehow compensated by S (mean over all R/S values = 0.98, standard deviation = 0.16). However, differences in the ratio existed across brain regions. This is illustrated in figure 4, where values of the ratio R/S bigger than 1 indicate supra-unbalanced areas, whereas smaller than 1 indicate infra-unbalanced areas. Panel a shows the highest R/S value corresponding to modules 9 and 10, which forms the default mode network. Similar to what happens for R and S, the ratio R/S is dependent on age (oneway ANOVA, p= 4e^−18^). Moreover, unlike to what happens for R, panel b shows the variations of R/S to be highly heteregeneous with age along lifespan (one-way anova, I1 vs I3 p= 0.001, I1 vs I4 p= 1e^−6^, I2 vs I3 p= 0.003, I2 vs I4 p= 1e^−9^ and I3 vs I4 p= 2e^−15^).

**Figure 4:**
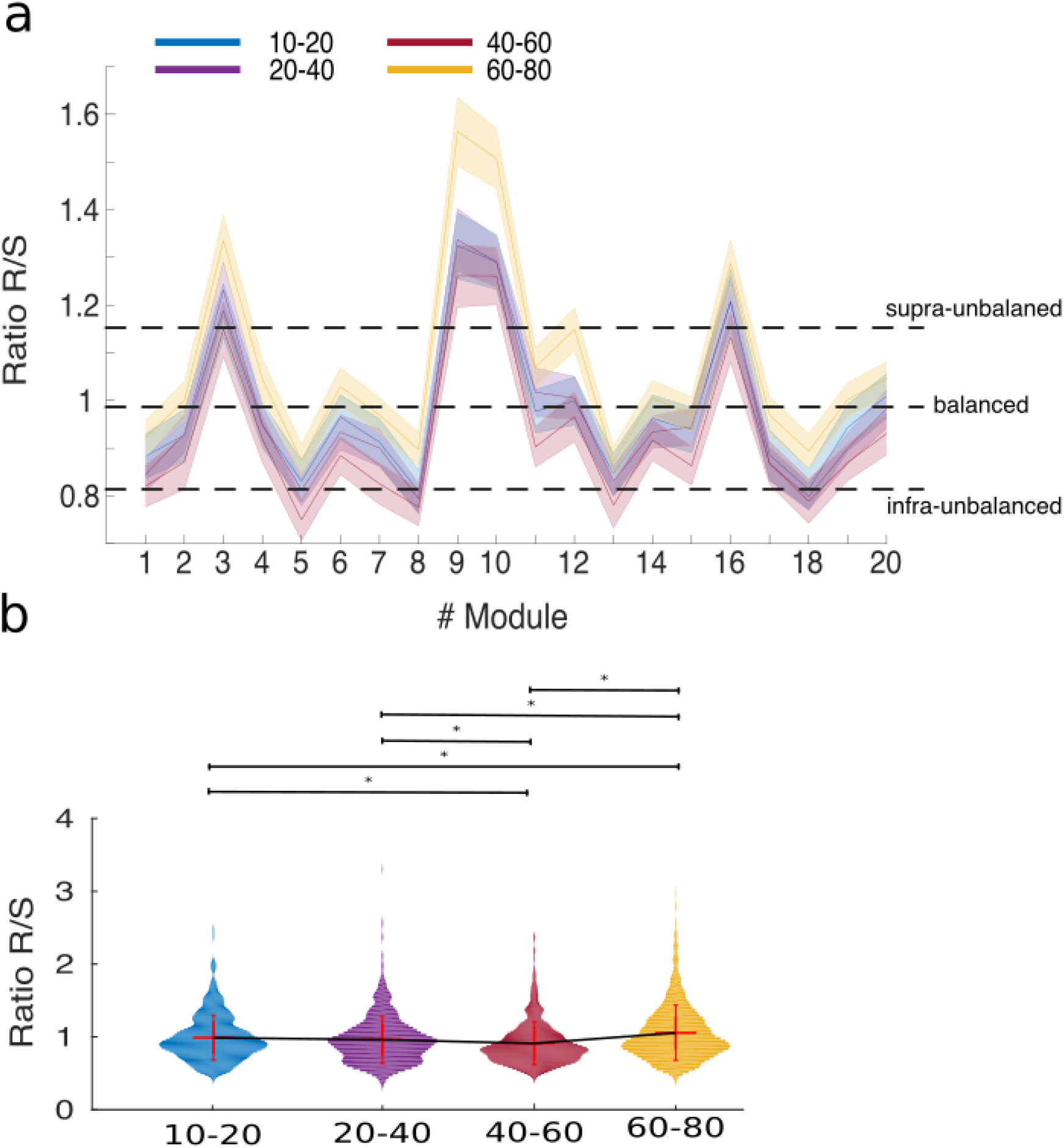
Across subjects, ratio R/S per brain module and along lifespan. **a:** Similar to figures 2 and 3, values are represented in 4 different age ranges, blue (10-20 years old), purple (20-40), red (40-60) and magenta (60-80). Three dashed lines delimite the three regimes: infra-unbalanced and supra-unbalanced, respectively, with values of R/S being smaller or bigger than mean minus or plus one times the standard deviation, and balanced, elsewhere. Dark central lines represent average values across subjects and shaded areas represent statistical error. Modules 9 and 10 corresponding to the default mode network are highly supra-unbalanced. Modules 5 and 8 corresponding to the fronto-parietal network are infra-unbalanced. **b:** Violin plots of R/S averaging over all modules and subjects within age interval. Modules * represents statistical significant differences after Bonferroni correction.

Next, we defined brain maps of infra-unbalanced R/S by looking to the brain areas with ratio smaller than the mean minus one times the standard deviation. Similarly, supra-unbalanced brain maps were determined by looking to the values bigger than the mean plus one times the standard deviation. Balanced areas corresponded to all the other values. Table 1 shows the overlapping of the three classes of brain maps (infra-unbalanced, supra-unbalanced and balanced) with the most important resting state networks: default mode, cerebellum, executive control, frontoparietal, sensorimotor and visual. Very remarkably, infra-unbalanced brain maps overlapped 9.5% with the frontoparietal network. Balanced brain maps overlapped 84% and 77% respectively with the cerebellum and visual networks. Supra-unbalanced brain maps matched 69.18% with the default mode network.

**Table 1:** Overlapping percentage of the ratio R/S with the main Resting State Networks

|  | Infra-balanced (%) | Balanced (%) | Supra-balanced (%) |
| --- | --- | --- | --- |
| **DMN** | 0.2246 | 19.0583 | 69.1893 |
| **Auditory** | 0.0284 | 38.1631 | 55.2979 |
| **Cerebellum** | 0 | 84.3334 | 0.8268 |
| **Executive Control** | 5.176 | 37.3482 | 30.9205 |
| **Frontoparietal** | 9.5896 | 39.358 | 39.5368 |
| **Sensorimotor** | 1.72 | 11.7076 | 57.8549 |
| **Visual** | 0 | 77.7678 | 5.9748 |

## Discussion

Interaction information (II) allows to assess how information between pairs of variables is enhanced (by synergy, S) or ignored (by redundancy, R) after adding a new interacting variable. Indeed, it is precisely the sign of II the one that reveals S (negative II) or R (positive II).

Here, we have studied how the values of R and S are distributed across brain areas and along lifespan. Brain areas with highest value of S were found subcortically in amygdala, hippocampus, putamen and caudate, although cortically as well in the entorhinal cortex, fusiform and temporal pole. With regard to R, we have found highly association with the default mode network, a well-known resting state network shown to be altered in many different brain disorders. This, together with the fact that R is more invariant along lifespan as compared to S, suggests a new role for the default mode network, as an integrator of information achieved by increasing redundancy. Perhaps, the significant increase of R occurring for the old population suggests a physiological alteration of this redundant role of the default mode network when we age, known to be altered [37], but this issue needs a further clarification.

R and S seem to be balanced, suggesting compensatory informational mechanisms in brain networks. We have seen that the frontoparietal network, classically associated to attentional control [38], is the network most affected by infra-unbalanced ratio R/S, revealing a new synergetic role of this network from an informational perspective. Moreover, cerebellum and visual are the two networks most balanced, again revealing new informational roles for these networks. Finally, the default mode network is the one with highest supra-unbalanced ratio R/S, and therefore, confirming its highly redundant role.

About what might be the possible mechanisms underlying R and S in the brain, future studies should address if network topological metrics, such as integration or segregation are somehow related to synergy and redundancy, although this is far the scope of the present work.

## Acknowledgment

ID acknowledges financial support from Department of Education of the Basque Country, postdoctoral program. JR acknowledges financial support from the Minister of Education, Language Policy and Culture (Basque Government) under Doctoral Research Staff Improvement Programme. Paolo Bonifazi is funded by Ministerio Economia, Industria y Competitividad, Spain (grant no. SAF2015-69484-R). Stephan P. Swinnen acknowledges financial support from Research Foundation Flanders (grants no. G0898.18N and G0708.14N) and Excellence of Science (MEMODYN, 30446199), KU Leuven Special Research Fund (grant no. C16/15/070). JMC acknowledges financial support from Department of Economical Development and Infrastructure of the Basque Country, Elkartek Program (grant no. KK-2018/00032) and Ministerio Economia, Industria y Competitividad, Spain and FEDER (grant no. DPI2016-79874-R). JMC and PB are funded by Ikerbasque.

## References

[1] McGill, W.J. Multivariate Information Transmission. Psychometrika 1954, 19, 97–116. doi:10.1007/BF02289159.

[2] Erramuzpe, A.; Ortega, G.; Pastor, J.; de Sola, R.; Marinazzo, D.; Stramaglia, S.; Cortes, J. Identification of redundant and synergetic circuits in triplets of electrophysiological data. J Neural Eng 2015, 12, 066007. doi:10.1088/1741-2560/12/6/066007.

[3] Lizier, J.T.; Atay, F.M.; Jost, J. Information Storage, Loop Motifs, and Clustered Structure in Complex Networks. Physical Review E 2012, 86. doi:10.1103/PhysRevE.86.026110.

[4] Wibral, M.; Lizier, J.T.; Vogler, S.; Priesemann, V.; Galuske, R. Local Active Information Storage as a Tool to Understand Distributed Neural Information Processing. Frontiers in Neuroinformatics 2014, 8. doi:10.3389/fninf.2014.00001.

[5] Conger, A.J. A Revised Definition for Suppressor Variables: A Guide To Their Identification and Interpretation. Educational and Psychological Measurement 1974, 34, 35–46. doi:10.1177/001316447403400105.

[6] Antonov, A.V.; Tetko, I.V.; Mader, M.T.; Budczies, J.; Mewes, H.W. Optimization Models for Cancer Classification: Extracting Gene Interaction Information from Microarray Expression Data. Bioinformatics 2004, 20, 644–652. doi:10.1093/bioinformatics/btg462.

[7] Wang, K.; Saito, M.; Bisikirska, B.C.; Alvarez, M.J.; Lim, W.K.; Rajbhandari, P.; Shen, Q.; Nemenman, I.; Basso, K.; Margolin, A.A.; Klein, U.; Dalla-Favera, R.; Califano, A. Genome-Wide Identification of Post-Translational Modulators of Transcription Factor Activity in Human B Cells. Nature Biotechnology 2009, 27, 829–837. doi:10.1038/nbt.1563.

[8] Marinazzo, D.; Gosseries, O.; Boly, M.; Ledoux, D.; Rosanova, M.; Massimini, M.; Noirhomme, Q.; Laureys, S. Directed information transfer in scalp electroencephalographic recordings: insights on disorders of consciousness. Clin EEG Neurosci 2014, 45, 33–39. doi:doi:10.1177/1550059413510703.

[9] Stramaglia, S.; Angelini, L.; Wu, G.; Cortes, J.M.; Faes, L.; Marinazzo, D. Synergetic and Redundant Information Flow Detected by Unnormalized Granger Causality: Application to Resting State fMRI. IEEE Transactions on Biomedical Engineering 2016, 63, 2518–2524. doi:10.1109/TBME.2016.2559578.

[10] Boveroux, P.; Vanhaudenhuyse, A.; Bruno, M.A.; Noirhomme, Q.; Lauwick, S.; Luxen, A.; Degueldre, C.; Plenevaux, A.; Schnakers, C.; Phillips, C.; Brichant, J.F.; Bonhomme, V.; Maquet, P.; Greicius, M.D.; Laureys, S.; Boly, M. Breakdown of Within-and between-Network Resting State Functional Magnetic Resonance Imaging Connectivity during Propofol-Induced Loss of Consciousness:. Anesthesiology 2010, 113, 1038–1053. doi:10.1097/ALN.0b013e3181f697f5.

[11] Noirhomme, Q.; Soddu, A.; Lehembre, R.; Vanhaudenhuyse, A.; Boveroux, P.; Boly, M.; Laureys, S. Brain Connectivity in Pathological and Pharmacological Coma. Frontiers in Systems Neuroscience 2010, 4. doi:10.3389/fnsys.2010.00160.

[12] Heine, L.; Soddu, A.; Gómez, F.; Vanhaudenhuyse, A.; Tshibanda, L.; Thonnard, M.; Charland-Verville, V.; Kirsch, M.; Laureys, S.; Demertzi, A. Resting State Networks and Consciousness. Frontiers in Psychology 2012, 3. doi:10.3389/fpsyg.2012.00295.

[13] Maki-Marttunen, V.; Diez, I.; Cortes, J.M.; Chialvo, D.R.; Villarreal, M. Disruption of Transfer Entropy and Inter-Hemispheric Brain Functional Connectivity in Patients with Disorder of Consciousness. Frontiers in Neuroinformatics 2013, 7. doi:10.3389/fninf.2013.00024.

[14] Woodward, N.D.; Rogers, B.; Heckers, S. Functional Resting-State Networks Are Differentially Affected in Schizophrenia. Schizophrenia Research 2011, 130, 86–93. doi:10.1016/j.schres.2011.03.010.

[15] Karbasforoushan, H.; Woodward, N.D. Resting-State Networks in Schizophrenia. Current Topics in Medicinal Chemistry 2012, 12, 2404–2414.

[16] Liao, W.; Zhang, Z.; Pan, Z.; Mantini, D.; Ding, J.; Duan, X.; Luo, C.; Lu, G.; Chen, H. Altered Functional Connectivity and Small-World in Mesial Temporal Lobe Epilepsy. PLoS ONE 2010, 5, e8525. doi:10.1371/journal.pone.0008525.

[17] Li, S.J.; Li, Z.; Wu, G.; Zhang, M.J.; Franczak, M.; Antuono, P.G. Alzheimer Disease: Evaluation of a Functional MR Imaging Index as a Marker. Radiology 2002, 225, 253–259. doi:10.1148/radiol.2251011301.

[18] Greicius, M.D.; Srivastava, G.; Reiss, A.L.; Menon, V. Default-Mode Network Activity Distinguishes Alzheimer’s Disease from Healthy Aging: Evidence from Functional MRI. Proceedings of the National Academy of Sciences 2004, 101, 4637–4642. doi:10.1073/pnas.0308627101.

[19] Rombouts, S.A.; Barkhof, F.; Goekoop, R.; Stam, C.J.; Scheltens, P. Altered Resting State Networks in Mild Cognitive Impairment and Mild Alzheimer’s Disease: An fMRI Study. Human Brain Mapping 2005, 26, 231–239. doi:10.1002/hbm.20160.

[20] Binnewijzend, M.A.; Schoonheim, M.M.; Sanz-Arigita, E.; Wink, A.M.; van der Flier, W.M.; Tolboom, N.; Adriaanse, S.M.; Damoiseaux, J.S.; Scheltens, P.; van Berckel, B.N.; Barkhof, F. Resting-State fMRI Changes in Alzheimer’s Disease and Mild Cognitive Impairment. Neurobiology of Aging 2012, 33, 2018–2028. doi:10.1016/j.neurobiolaging.2011.07.003.

[21] Sheline, Y.I.; Raichle, M.E. Resting State Functional Connectivity in Preclinical Alzheimer’s Disease. Biological Psychiatry 2013, 74, 340–347. doi:10.1016/j.biopsych.2012.11.028.

[22] Bonifazi, P.; Erramuzpe, A.; Diez, I.; Gabilondo, I.; Boisgontier, M.; Pauwels, L.; Stramaglia, S.; Swinnen, S.; Cortes, J. Structure-function multi-scale connectomics reveals a major role of the fronto-striato-thalamic circuit in brain aging. Hum Brain Mapp 2018. doi:10.1002/hbm.24312.

[23] Marinazzo, D.; Pellicoro, M.; Wu, G.; Angelini, L.; Cortes, J.M.; Stramaglia, S. Information Transfer and Criticality in the Ising Model on the Human Connectome. PLoS ONE 2014, 9, e93616. doi:10.1371/journal.pone.0093616.

[24] Alonso-Montes, C.; Diez, I.; Remaki, L.; Escudero, I.n.; Mateos, B.; Rosseel, Y.; Marinazzo, D.; Stramaglia, S.; Cortes, J.M. Lagged and Instantaneous Dynamical Influences Related to Brain Structural Connectivity. Frontiers in Psychology 2015, 6. doi:10.3389/fpsyg.2015.01024.

[25] Amor, T.A.; Russo, R.; Diez, I.; Bharath, P.; Zirovich, M.; Stramaglia, S.; Cortes, J.M.; de Arcangelis, L.; Chialvo, D.R. Extreme Brain Events: Higher-Order Statistics of Brain Resting Activity and Its Relation with Structural Connectivity. EPL (Europhysics Letters) 2015, 111, 68007. doi:10.1209/0295-5075/111/68007.

[26] Diez, I.; Erramuzpe, A.; Escudero, I.n.; Mateos, B.; Cabrera, A.; Marinazzo, D.; Sanz-Arigita, E.J.; Stramaglia, S.; Cortes Diaz, J.M.; for the Alzheimer’s Disease Neuroimaging Initiative. Information Flow Between Resting-State Networks. Brain Connectivity 2015, 5, 554–564. doi:10.1089/brain.2014.0337.

[27] Rasero, J.; Pellicoro, M.; Angelini, L.; Cortes, J.M.; Marinazzo, D.; Stramaglia, S. Consensus Clustering Approach to Group Brain Connectivity Matrices. Network Neuroscience 2017, 1, 242–253. doi:10.1162/NETN_a_00017.

[28] Stramaglia, S.; Pellicoro, M.; Angelini, L.; Amico, E.; Aerts, H.; Cortes, J.M.; Laureys, S.; Marinazzo, D. Ising Model with Conserved Magnetization on the Human Connectome: Implications on the Relation Structure-Function in Wakefulness and Anesthesia. Chaos: An Interdisciplinary Journal of Nonlinear Science 2017, 27, 047407. doi:10.1063/1.4978999.

[29] Cordes, D.; Haughton, V.M.; Arfanakis, K.; Carew, J.D.; Turski, P.A.; Moritz, C.H.; Quigley, M.A.; Meyerand, M.E. Frequencies Contributing to Functional Connectivity in the Cerebral Cortex in “Resting-State” Data. AJNR. American journal of neuroradiology 2001, 22, 1326–1333.

[30] Yan, C.G.; Cheung, B.; Kelly, C.; Colcombe, S.; Craddock, R.C.; Di Martino, A.; Li, Q.; Zuo, X.N.; Castellanos, F.X.; Milham, M.P. A Comprehensive Assessment of Regional Variation in the Impact of Head Micromovements on Functional Connectomics. NeuroImage 2013, 76, 183–201. doi:10.1016/j.neuroimage.2013.03.004.

[31] Diez, I.; Bonifazi, P.; Escudero, I.n.; Mateos, B.; Muñoz, M.A.; Stramaglia, S.; Cortes, J.M. A Novel Brain Partition Highlights the Modular Skeleton Shared by Structure and Function. Scientific Reports 2015, 5. doi:10.1038/srep10532.

[32] Diez, I.; Drijkoningen, D.; Stramaglia, S.; Bonifazi, P.; Marinazzo, D.; Gooijers, J.; Swinnen, S.P.; Cortes, J.M. Enhanced Prefrontal Functional-Structural Networks to Support Postural Control Deficits after Traumatic Brain Injury in a Pediatric Population. Network Neuroscience 2017, 1, 116–142. doi:10.1162/NETN_a_00007.

[33] Jaynes, E.T. Information Theory and Statistical Mechanics. Physical Review 1957, 106, 620–630. doi:10.1103/PhysRev.106.620.

[34] Cover, T.M.; Thomas, J.A. Elements of Information Theory, 2nd ed ed.; Wiley-Interscience: Hoboken, N.J, 2006. OCLC: ocm59879802.

[35] Ince, R.; Giordano, B.; Kayser, C.; Rousselet, G.; Gross, J.; Schyns, P. A statistical framework for neuroimaging data analysis based on mutual information estimated via a Gaussian copula. Hum Brain Mapp 2017, 38, 1541–1573. doi:10.1101/043745.

[36] Smith, S.; Fox, P.; Miller, K.; Glahn, D.; Fox, P.; Mackay, C.; Filippini, N.; Watkins, K.; Toro, R.; Laird, A.; Beckmann, C. Correspondence of the brain’s functional architecture during activation and rest. Proc Natl Acad Sci USA 2009, 106, 535816. doi:10.1073/pnas.0905267106.

[37] Mevel, K.; Chetelat, G.; Eustache, F.; Desgranges, B. The default mode network in healthy aging and Alzheimer’s disease. Int J Alzheimers Dis 2011, 2011, 13040–13045. doi:10.4061/2011/535816.

[38] Scolari, M.; Seidl-Rathkopf, K.; Kastner, S. Functions of the human frontoparietal attention network: Evidence from neuroimaging. Curr Opin Behav Sci 2015, 1, 32–39. doi:10.1016/j.cobeha.2014.08.003.

